# Phosphorylation of Elovl5 changes its substrate preference to synthesize Mead acid in response to essential fatty acid deficiency

**DOI:** 10.1101/2020.01.31.929224

**Authors:** Yuri Hayashi, Misato Yamano, Nozomu Kono, Hiroyuki Arai, Yoko Fujiwara, Ikuyo Ichi

**Affiliations:** Graduate School of Humanities and Sciences, Ochanomizu University, Tokyo, Japan; Graduate School of Pharmaceutical Sciences, University of Tokyo, Tokyo, Japan; Graduate School of Medicine, University of Tokyo, Tokyo, Japan; Natural Science Division, Faculty of Core Research, Ochanomizu University, Tokyo, Japan; Institute for Human Life Innovation, Ochanomizu University, Tokyo, Japan

**Keywords:** polyunsaturated fatty acid (PUFA), essential fatty acid (EFA), Elovl5, fatty acid metabolism, phosphorylation, glycogen synthase kinase 3 (GSK-3), membrane protein

## Abstract

Polyunsaturated fatty acids (PUFAs) of the n-6 and n-3 series cannot be synthesized in mammals and therefore are called essential fatty acids (EFAs). Mead acid (20:3n-9) is an unusual n-9 PUFA, endogenously synthesized from oleic acid (18:1n-9) in an EFA-deficient state. Although Elovl5, a fatty acid elongase, has long been known to selectively elongate C18 and C20 PUFAs, it can use 18:1n-9 as a substrate for the synthesis of Mead acid under C20 PUFA-deficient, but not-sufficient, conditions. We found, by an in vitro enzyme assay, that the microsomal fraction obtained from PUFA-deficient, but not -sufficient, cells showed significant Elovl5 activity toward 18:1n-9, with no effect on its constitutive activity toward 18:3n-6, implying that Elovl5 acquires the activity toward 18:1n-9 under the PUFA-deficient conditions at the enzyme level. Further biochemical analysis revealed that Elovl5 was phosphorylated in the C20 PUFA-supplemented cells, and that treatment with an inhibitor of glycogen synthase kinase 3 (GSK3) completely abolished the phosphorylation of Elovl5 and retained the Elovl5 activity toward 18:1n-9, even in the presence of C20 PUFA. Finally, mutation of putative phosphorylation sites (T281A/S283A/S285A) on Elovl5 did not decrease the activity of Elovl5 toward 18:1n-9 by supplementation with C20 PUFA, suggesting that the phosphorylation of Elovl5 contributed to a change in substrate preference. Thus, by changing its substrate specificity in an EFA-deficient state, Elovl5 is able to regulate the synthesis of Mead acid to maintain levels of long-chain PUFAs.

---

Polyunsaturated fatty acids (PUFAs) are important components of biological membranes and the precursors of bioactive substances such as prostaglandins, which mediate numerous physiological and pathological processes (1). Mammals are unable to synthesize n-6 or n-3 series PUFAs and must obtain them from diets. Therefore, PUFAs such as linoleic acid (18:2n-6) and α-linolenic acid (18:3n-3) are called essential fatty acids (EFAs), which can be converted to long-chain and highly unsaturated fatty acids, such as arachidonic acid (20:4n-6) and eicosapentaenoic acid (EPA, 20:5n-3), through elongation and desaturation reactions (Fig. 1A). EFA-deficiency causes impaired growth, skin lesions, and infertility in rodents (2, 3), and occurs in humans when the contents of EFAs in diets are less than 1–2% of the total calories (5). Indeed, EFA-deficiency is observed in patients with intestinal lipid malabsorption and in patients that receive total parenteral nutrition with fat restriction (6, 7).

**Figure 1.**
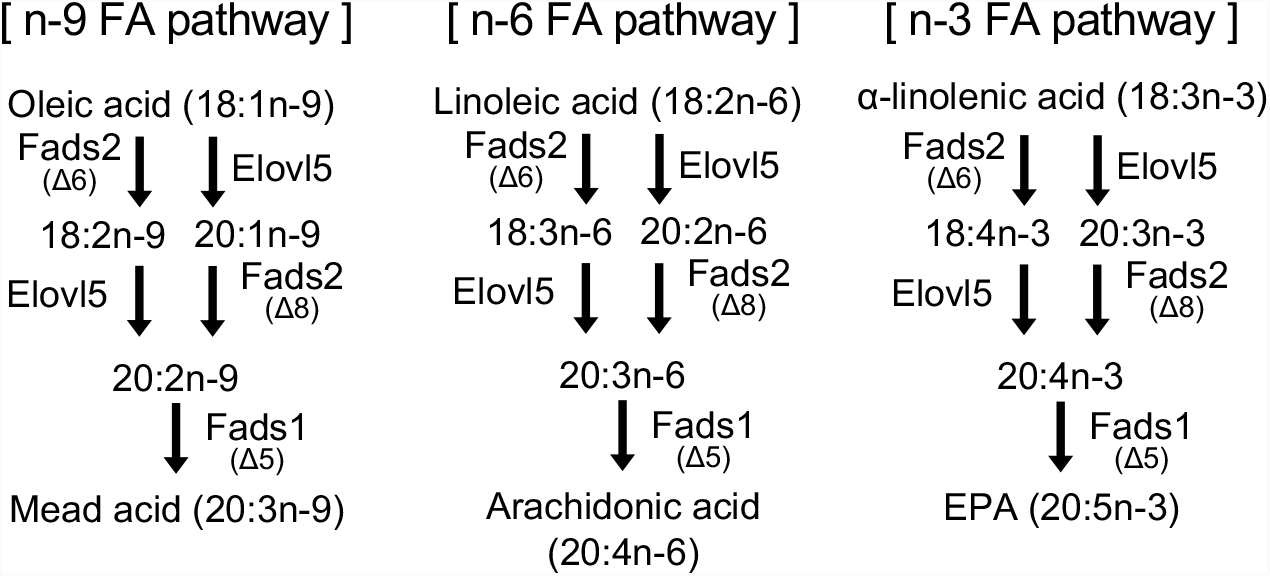
Synthetic pathways of mammal PUFAs.

In the EFA-deficient state, 5,8,11-eicosatrienoic acid (Mead acid, 20:3n-9) is endogenously synthesized from 18:1n-9 (oleic acid) (Fig. 1A) (4), and plasma and tissue levels of Mead acid are used as a diagnostic indicator of EFA-deficiency (2, 3). Among infants in intensive care units, those with a high level of Mead acid grew more slowly than other infants despite receiving similar amounts of macronutrients such as proteins and carbohydrates (8). Mead acid is sometimes detected in human cancers such as breast cancer (9) and prostate cancer (10), suggesting that some cancer tissues are in the EFA-deficient state.

Mead acid is thought to be used as a substitute for essential PUFAs in biological membranes. For example, incorporation of PUFAs such as arachidonic acid (20:4n-6) into phospholipids is required for the efficient transport of dietary lipids in the intestine and the liver (11, 12), and Mead acid in membrane phospholipids can rescue the impaired hepatic lipid transport in the EFA-deficient state (13). Oxidative metabolites of Mead acid participate in both proinflammatory (14) and anti-inflammatory signaling pathways (15), suggesting that Mead acid at least partly compensates for the reduced levels of n-6 and n-3 PUFAs. Mead acid also exerts anti-angiogenic activity to attenuate the activity of osteoblasts (16, 17) and to inhibit tumorigenesis in certain types of cancer (18). Moreover, intraperitoneal injection of Mead acid ameliorates allergic skin inflammation in the murine contact hypersensitivity model (19). Elucidation of Mead acid synthesis and metabolism may therefore contribute to the fields of immunology and cancer as well as to development of a novel strategy to treat patients with EFA-deficiency. We previously showed that Mead acid is produced from oleic acid (18:1n-9) by two desaturation enzymes (Fads1 and Fads2) and one elongation enzyme (Elovl5) (20) (Fig. 1A). These three enzymes also participate in the synthesis of 20:4n-6 from 18:2n-6 and the synthesis of 20:5n-3 from 18:3n-3 (Fig. 1A).

Elovl, a transmembrane protein residing in the endoplasmic reticulum (ER), is one of group of enzymes that control the rate of elongation of fatty acids with a chain length of more than 16 carbon atoms. The Elovl family is composed of seven distinct subtypes (Elovl1-7) in mammals, and each Elovl enzyme has a defined substrate specificity depending on the chain length or degree of unsaturation of fatty acids (21). However, there is a discrepancy regarding the substrate specificity of Elovl5. While Elovl5 generally elongates C18 and C20 PUFAs in mammals (22, 23), we have demonstrated that Elovl5 had activity toward the C18 monounsaturated fatty acid 18:1n-9 in an elongation assay using the microsomal fraction obtained from cells in an EFA-deficient state, and that this elongation activity was involved in the production of Mead acid in the cells (20). Moreover, there was a dramatic increase of 20:1n-9 in tissues of animals fed an EFA-deficient diet (20), supporting the conversion of 18:1n-9 to 20:1n-9 *in vivo* (Fig. 1A). However, considering that high levels of 18:1n-9 are still maintained under EFA-sufficient conditions, it remains unknown why 18:1n-9 cannot serve as a substrate for Elovl5 in the presence of PUFAs. Although this could be simply because Elovl5 has a higher substrate specificity for C18 and C20 PUFAs than for 18:1n-9, the existence of an alternative mechanism that influences Elovl5 activity cannot be ruled out.

In this study, we compared the substrate specificities of Elovl5 under EFA-sufficient and - deficient conditions and found that, in the EFA-deficient state, Elovl5 acquires an enzyme activity towards 18:1n-9, leading to the synthesis of Mead acid (20:3n-9), without affecting the activity towards 18:3n-6. Our results also suggest that the phosphorylation of Elovl5 causes a change in the substrate specificity of Elovl5 between EFA-sufficient and -deficient states. Thus, we found a novel regulatory mechanism in which Elovl5 changes the substrate preference by a post-translational modification.

## Results

### The elongation products of Elovl5 in n-9 and n-6 fatty acid pathways are differentially controlled by PUFAs

We previously showed that NIH3T3 cells cultured in a medium containing 10% (v/v) fetal bovine serum (FBS) produced significant amounts of Mead acid, a marker fatty acid of EFA-deficiency (20). In fact, the PUFA level in 10% FBS is much lower than that in mouse plasma (Fig. S1), upon addition of PUFAs [25 μM arachidonic acid (20:4n-6) + 25 μM EPA (20:5n-3)] to the culture medium, the level of Mead acid was greatly decreased (Fig. 2A and B). Treatment with PUFAs also reduced the intermediate products (18:2n-9, 20:1n-9, and 20:2n-9) for the synthesis of Mead acid in the n-9 fatty acid (FA) pathway (Fig. 2A and B). Furthermore, the levels of Mead acid and its intermediate products were decreased to the same degree in NIH3T3 cells supplemented with n-3 PUFAs [25 μM α-linolenic acid (18:3n-3) + 25 μM EPA (20:5n-3)] or with n-6 PUFAs [25 μM linoleic acid (18:2n-6) + 25 μM arachidonic acid (20:4n-6)] (Fig. S2), suggesting that the synthesis of 18:1n-9 from Mead acid was similarly suppressed by the supplemented n-3 and n-6 PUFAs.

**Figure 2.**
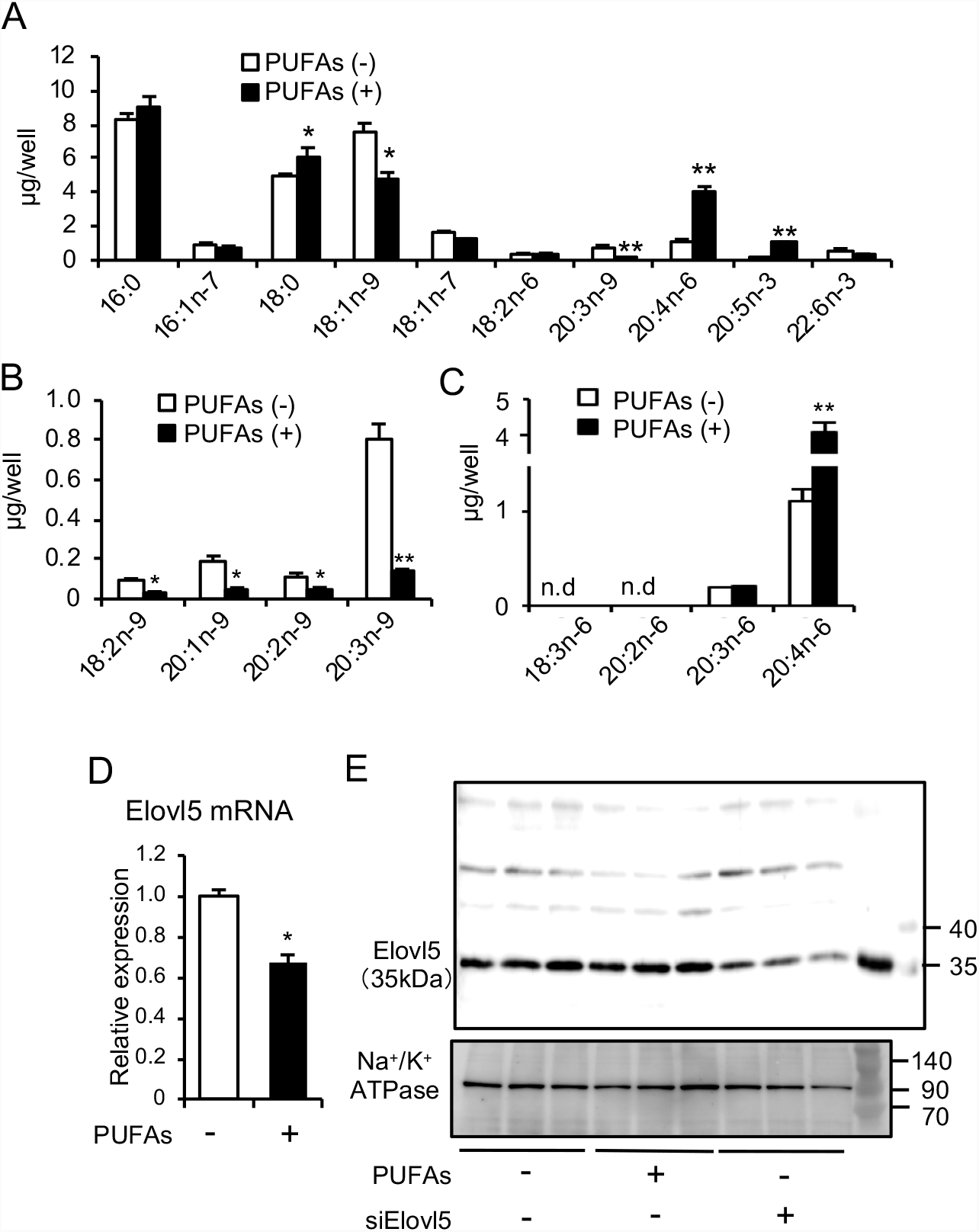
Change of fatty acid composition with addition of PUFAs. (A) Change of fatty acid composition in NIH3T3 cells with or without addition of the PUFAs [25 μM arachidonic acid (20:4n-6) and 25 μM EPA (20:5n-3)] for 24 h. Asterisks indicate statistically significant differences compared to PUFAs (-) (*p < 0.05, **p < 0.01 by Student’s t-test). (B, C) Change of the intermediate products of n-9 PUFAs (B) and n-6 PUFAs (C) with or without addition of the PUFAs for 24 h. Asterisks indicate statistically significant differences compared to PUFAs (-) (*p < 0.05, **p < 0.01 by Student’s t-test). (D) mRNA expression of Elovl5 with or without addition of PUFAs in NIH3T3 cells. Asterisks indicate statistically significant differences compared to PUFAs (-) (*p < 0.05 by Student’s t-test). (E) The Elovl5 proteins in the NIH3T3 cells treated with or without the PUFAs and in the cells treated with siElovl5 were determined by immunoblotting with Elovl5 antibody.

Adding PUFAs decreased the cellular levels of both 18:1n-9 and 20:1n-9 (Fig. 2A and B). The level of 20:1n-9 was not increased by adding 18:1n-9 to the PUFA-treated cells (Fig. S3A and B). Thus, the decrease of 20:1n-9 was not due to the reduction of 18:1n-9, a direct precursor for 20:1n-9. It is more likely that it was caused by the reduced elongation activity of Elovl5 from 18:1n-9 to 20:1n-9. On the other hand, the level of 20:3n-6 was not affected by the addition of PUFAs (Fig. 2C). 20:3n-6 is produced through the elongation of 18:2n-6 and 18:3n-6 by Elovl5 (Fig. 1A). Although 18:3n-6 and 20:2n-6 were nearly undetectable in NIH3T3 cells under the experimental conditions used, the above result suggests that the elongation activity of Elovl5 in the n-6 FA pathway was not altered by PUFAs (see Fig. 1A). Taken together, these results raised the possibility that the Elovl5 activities toward n-9 and n-6 FAs are differentially controlled by PUFAs.

The addition of PUFAs to NIH3T3 cells decreased the mRNA level of Elovl5 by about 30% (Fig. 2D), which is consistent with the previous observation that PUFAs suppress the hepatic expression of Elovl5 mRNA via inactivation of the LXRα-SREBP1c pathway (24). However, western blots revealed that PUFAs did not significantly change the protein expression of Elovl5 (Fig. 2E). Thus, it is unlikely that the decreased level of 20:1n-9 in the presence of PUFAs is due to the reduction of the Elovl5 protein level.

### Effect of PUFAs on Elovl5 activity toward 18:1n-9-CoA and 18:3n-6-CoA

We then evaluated the elongation activity of Elovl5 toward 18:1n-9-CoA and 18:3n-6-CoA with an *in vitro* enzyme assay, where the membrane fraction of NIH3T3 cells was incubated with 18:1n-9-CoA or 18:3n-6-CoA as an acceptor and malonyl-CoA as a donor, and then the product, 20:1n-9 or 20:3n-6, was measured with GC-MS after saponification (Fig. 3A). We chose 18:3n-6-CoA rather than 18:2n-6-CoA as a substrate for Elovl5 because the former was reported to be a better substrate for Elovl5 (23). 20:1n-9 and 20:3n-6 were increased following incubation with 18:1n-9-CoA and 18:3n-6-CoA, respectively (Fig. 3B and C), indicating that the elongation reaction against both substrates proceeded under the assay conditions used. Although several other elongases, including Elovl3, 6, and 7, have been reported to elongate 18:1n-9-CoA (23, 25), the elongation activity toward 18:1n-9 was markedly reduced by siRNA-mediated knockdown of Elovl5 (Fig. 3E), in which the expression of other Elovl isoforms were unaffected (Fig. 3D), confirming that Elovl5 made a major contribution to the production of 20:1n-9. Similarly, the elongation activity from 18:3n-6 to 20:3n-6 in the n-6 FA pathway was likely due to Elovl5 (Fig. 3F). In this pathway, the endogenous basal level of 20:3n-6 (white bars in Fig. 3F) was decreased in Elovl5 knockdown cells, possibly because the production of 20:3n-6 was suppressed during siElovl5 treatment.

**Figure 3.**
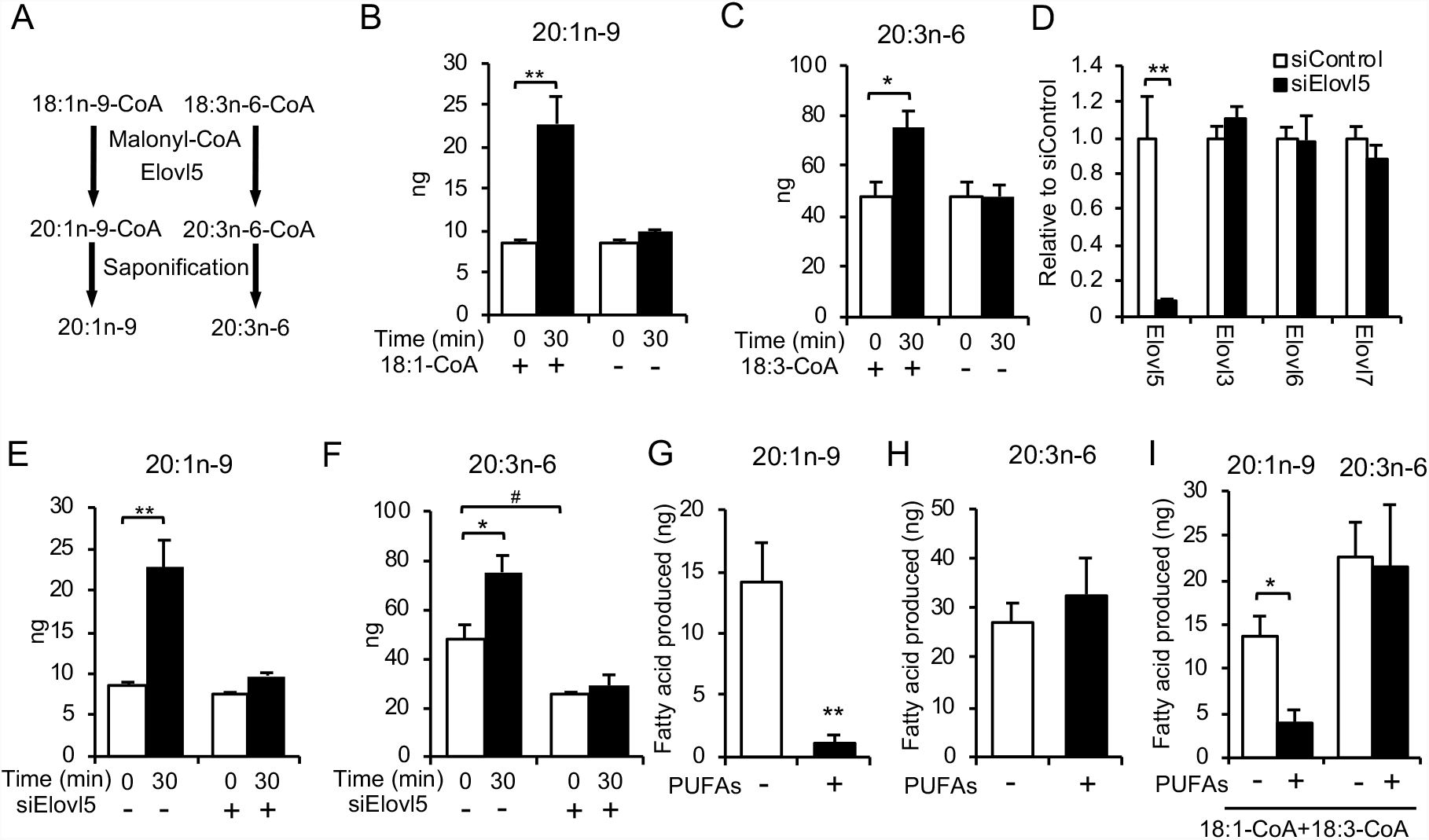
Effect of addition of PUFAs on Elovl5 activity toward 18:1n-9-CoA or 18:3n-6-CoA. (A) Schema of elongation assay of Elovl5 toward 18:1n-9-CoA and 18:3n-6-CoA. (B, C) Total membrane proteins in NIH3T3 cells were incubated with or without 50 μM 18:1n-9-CoA (B) or 18:3n-6-CoA (C) for 0 and 30 min. The amount of 20:1n-9 (B) or 20:3n-6 (C) was determined by GC-MS respectively. Statistically significant differences compared to 0 min (*p < 0.05, **p < 0.01 by Student’s t-test). (D) The mRNA expression of each Elovl was measured in the cells treated with siElovl5. Asterisks indicate statistically significant differences compared to siControl (*p < 0.05, **p < 0.01 by Student’s t-test). (E, F) The amount of 20:1n-9 (E) or 20:3n-6 (F) in the cells treated with or without siElovl5 was determined for 0 and 30 min. Asterisks indicate statistically significant differences compared to 0 min (*p < 0.05, ** p < 0.01, by Student’s t-test). Sharp indicates statistically significant differences compared to siControl (#p < 0.05, by Student’s t-test) (G, H) The elongation activity toward 18:1n-9-CoA (G) or 18:3n-6-CoA (H) in the membrane obtained from the PUFA [25 μM arachidonic acid (20:4n-6) and 25 μM EPA (20:5n-3)]-treated or non-treated cells. The increased amounts of 20:1n-9 (G) or 20:3n-6 (H) products during the reaction are indicated. Asterisks indicate statistically significant differences compared to PUFAs (-) (**p < 0.01 by Student’s t-test). (I) Substrate competition for Elovl5 elongation between 18:1n-9-CoA and 18:3n-6-CoA in the cells treated with PUFAs. *In vitro* FA elongation assay was performed in the presence of both substrates 25 μM 18:1n-9-CoA and 25 μM 18:3n-6-CoA. The increased amounts of each elongated product (20:1n-9 or 20:3n-6) are indicated. Asterisks indicate statistically significant differences (*p < 0.05, **p < 0.01 by Student’s t-test).

The Elovl5 activity toward 18:1n-9, as expressed by the increase of 20:1n-9 after *in vitro* incubation, was greatly reduced in NIH3T3 cells supplemented with the PUFAs (Fig. 3G). In contrast, the Elovl5 activity toward 18:3n-6, as expressed by the increase of 20:3n-6, was not affected in the PUFA-treated cells (Fig. 3H). Furthermore, even in the co-presence of 18:1n-9-CoA and 18:3n-6-CoA as substrates in the same assay, only the Elovl5 activity toward 18:1n-9-CoA was decreased in the membrane fraction obtained from the PUFA-treated cells (Fig. 3I). These results suggest that adding PUFAs to EFA-deficient cells induced a marked reduction of the Elovl5 activity toward 18:1n-9-CoA, but not toward 18:3n-6-CoA.

### Cellular amount of C20 PUFAs is related to the Elovl5 activity toward 18:1n-9-CoA

We tested whether the addition of C18 PUFAs, linoleic acid (18:2n-6) and α-linolenic acid (18:3n-3), also reduced the Elovl5 activity toward 18:1n-9. The Elovl5 activity in NIH3T3 cells treated with a mixture of 25 μM 18:2n-6 and 25 μM 18:3n-3 was decreased by only 70%, relative to more than 90% reduction in the cells treated with C20 PUFAs (Fig. 4A). Since C18 PUFAs could be converted to C20 PUFAs in the cell, we examined whether the reduced Elovl5 activity was due to the C18 PUFAs themselves or their elongation products, C20 PUFAs. In fact, treating the cells with C18 PUFAs increased the cellular content of C20 PUFAs (20:3n-9, 20:3n-6, 20:4n-6 and 20:5n-3) as well as C18 PUFAs (18:2n-6 and 18:3n-3) (Fig. 4B), whereas the addition of C20 PUFAs increased only the cellular content of C20 PUFAs and slightly decreased the content of C18 PUFAs. We then attempted to suppress the conversion of C18 PUFAs to C20 PUFAs in the cells treated with C18 PUFAs by adding TOFA (5-(tetradecyloxy)-2-furancarboxylic acid), an inhibitor of acetyl CoA carboxylase that is an essential enzyme for the production of malonyl-CoA (Fig. 4C). The increase of C20 PUFAs in the cells treated with C18 PUFAs was appreciably suppressed by TOFA treatment (Fig. 4D). Under the experimental conditions used, the suppression of the Elovl5 activity toward 18:1n-9 in the membrane fraction by C18 PUFAs was mostly cancelled by TOFA (Fig. 4E). Thus, the suppression of the Elovl5 activity toward 18:1n-9 by C18 PUFAs appeared to be due to C20 PUFAs produced from C18 PUFAs.

**Figure 4.**
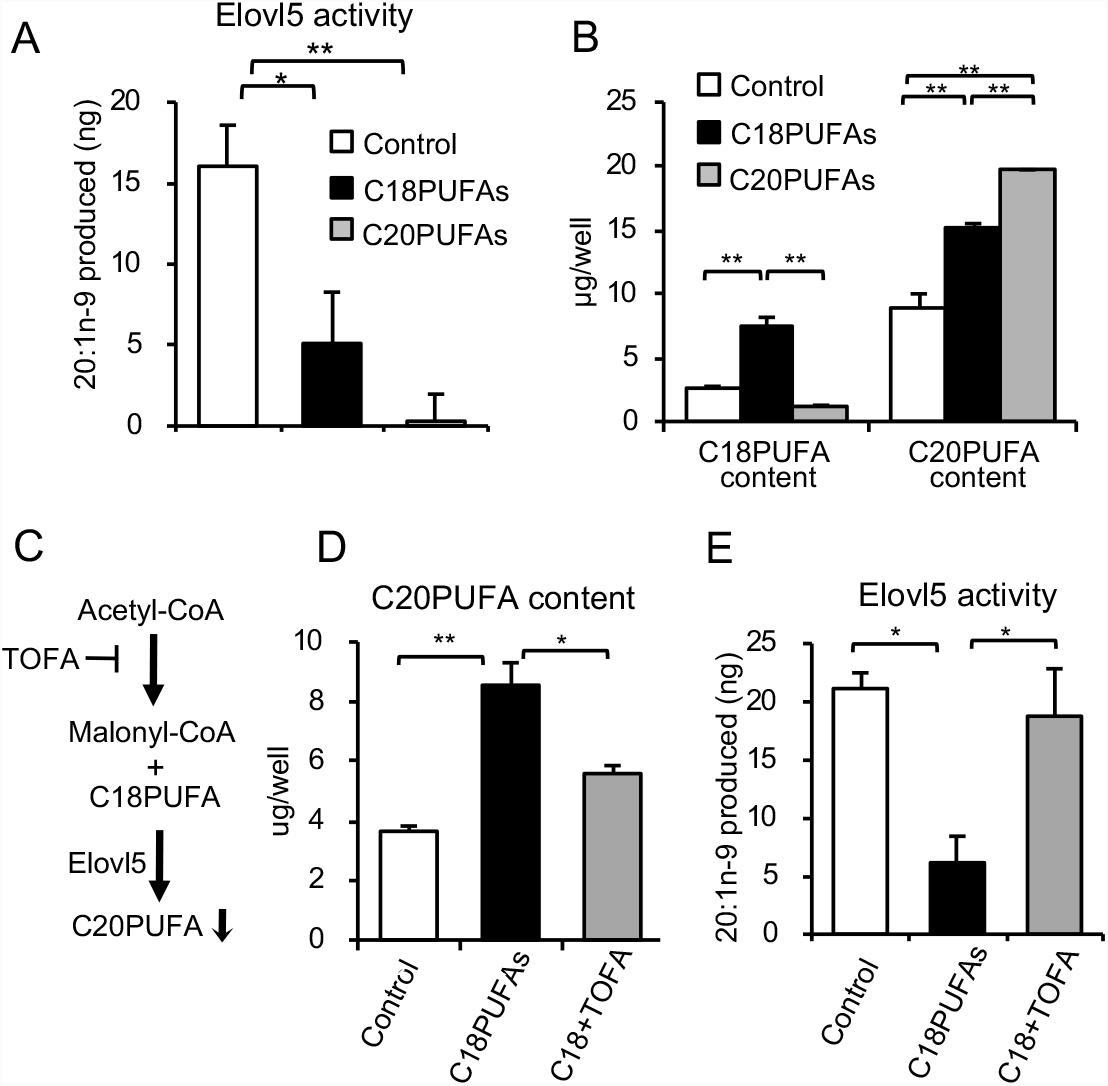
Regulation of the Elovl5 activity toward 18:1n-9 involves intracellular C20 PUFAs. (A) Elovl5 activity toward 18:1n-9-CoA was determined by *in vitro* FA elongation assay using membrane fractions from cells treated with C18 PUFAs (25 μM 18:2n-6 + 25 μM 18:3n-3) or C20 PUFAs (25 μM 20:4n-6 + 25 μM 20:5n-3). (B) The levels of C18PUFAs (18:2n-6 and 18:3n-3) and C20PUFAs (20:3n-9, 20:3n-6, 20:4n-6, and 20:5n-3) in the cells treated with C18 PUFAs (25 μM 18:2n-6 + 25 μM 18:3n-3) or with C20 PUFAs (25 μM 20:4n-6 + 25 μM 20:5n-3) were indicated. (C) Suppression of elongation of the C18 PUFAs in the cells treated with C18 PUFAs using acetyl-CoA carboxylase inhibitor, TOFA. (D) The levels of intracellular C20 PUFAs in the cells treated with C18 PUFAs and 10 μM TOFA. (E) Elovl5 activity toward 18:1n-9-CoA was determined by *in vitro* FA elongation assay from cells treated with C18 PUFAs and 10 μM TOFA. *p < 0.05, **p < 0.01 (Tukey-Kramer test).

### Elovl5 is phosphorylated in the presence of C20 PUFAs in a GSK3-dependent manner

The finding that C20 PUFAs greatly decreased Elovl5 activity toward 18:1n-9 indicated that the presence of C20 PUFAs changed the substrate specificity of Elovl5, although it did not change the Elovl5 activity toward 18:3n-6. We then investigated the possibility that post-translational modification of Elovl5 underlies its change in substrate specificity by C20 PUFAs. Elo2 is a key enzyme of very long chain fatty acid synthesis in yeast, in which only saturated fatty acids are thought to serve as substrates for Elo2. A recent study demonstrated that Elo2 was phosphorylated in nutrient-depleted cells (26), although it has not yet been established whether phosphorylation of Elo2 is important for the synthesis of very long chain fatty acids (VLCFAs) (27) or for the activity/stability of the enzyme (26). Phosphorylation sites are highly conserved between yeast Elo2 and mammalian Elovl5, but not other Elovl isoforms. Furthermore, a large-scale phosphorylation analysis revealed that human and mouse Elovl5 are phosphorylated on residues corresponding to the phosphorylated sites in yeast Elo2 (28, 29). We therefore examined whether the phosphorylation status of Elovl5 depends on the presence of C20 PUFAs in NIH3T3 cells. Phosphorylated proteins were separated using Phos-tag PAGE, and detected with an anti-Elovl5 antibody. Intriguingly, treatment of the cells with C20 PUFAs produced an up-shifted band of the Elovl5 protein (Fig. 5A), implying that the C20 PUFAs induced the phosphorylation of Elovl5.

**Figure 5.**
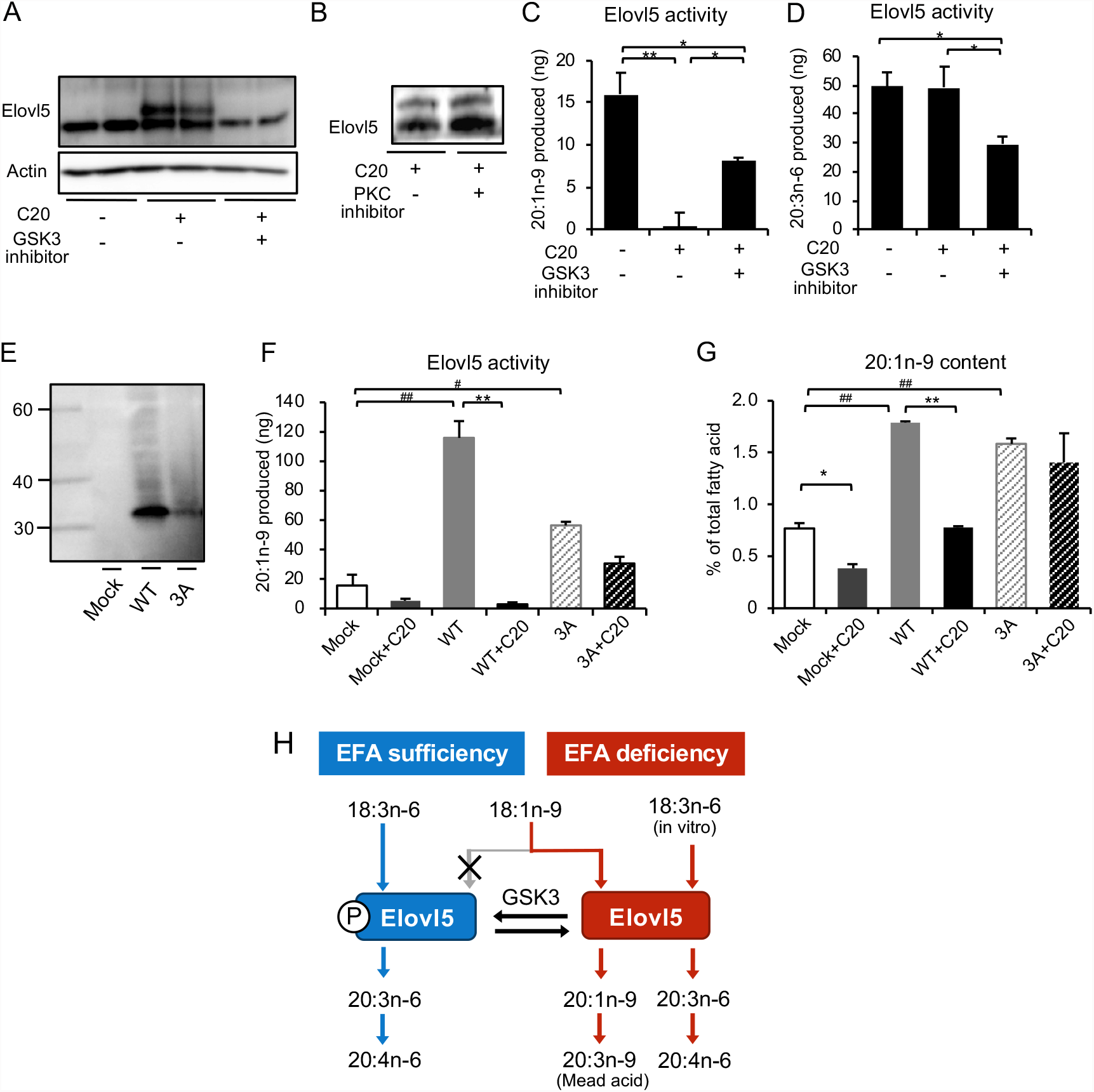
Regulation of Elovl5 activity toward 18:1n-9 by phosphorylation in the presence of C20 PUFAs. (A, B) NIH3T3 cells were treated with C20 PUFAs and two types of kinase inhibitors, GSK3 inhibitor (1 μM BIO) (A) and PKC inhibitor (1 μM BIM) (B) for 24 h. Phosphorylated Elovl5 was separated by Phos-tag PAGE, and was determined by immunoblotting with anti-Elovl5 antibody. (C, D) Elovl5 activities toward 18:1n-9-CoA (C) and 18:3n-6-CoA (D) were measured in cells that had been treated with BIO upon C20 PUFAs treatment. *p < 0.05, **p < 0.01 (Tukey-Kramer test). (E) N-terminal 3xFLAG-tagged wild-type (WT) or T281A/S283A/S285A triple mutant (3A) Elovl5 plasmids were transfected to HEK293T cells. FLAG-tagged Elovl5 was detected with anti-FLAG antibody. (F) Elovl5 activity toward 18:1n-9-CoA was determined by *in vitro* FA elongation assay from the cells treated with C20 PUFAs upon WT-Elovl5 or 3A-Elovl5 transfection. (G) Cellular amount of 20:1n-9 was determined by GC-MS in the cells treated with C20 PUFAs upon WT-Elovl5 or 3A-Elovl5 transfection. **p < 0.01 compared to PUFA (-), # p < 0.05, ## p < 0.01 compared to mock (Tukey-Kramer test). (H) Regulation of Elovl5 activity and Mead acid synthesis responding to EFA-deficiency.

The phosphorylation of Elo2 requires glycogen synthase kinase 3 (GSK3) (26) and occurs at sites that resemble GSK3 consensus sites (28). The mobility shift of Elovl5 by C20 PUFAs was completely abolished by the GSK3 inhibitor, 6-bromoindirubin-3’-oxime (BIO) (Fig. 4A), but not by the protein kinase C (PKC) inhibitor bisindolylmaleimide (BIM) (Fig. 5B), indicating that the phosphorylation of Elovl5 was also dependent on GSK3 in mammalian cells. Furthermore, BIO restored about half of the Elovl5 activity toward 18:1n-9 in the C20 PUFA-pretreated cells (Fig. 5C). The protein level of Elovl5 was decreased by about half in the cells treated with both C20 PUFAs and BIO (Fig. 5A). Although the reason was not clear, GSK3-dependent phosphorylation might be involved in the stabilization of Elovl5 protein. Taking into account of this fact, the GSK3 inhibitor appeared to nearly completely recover the Elovl5 activity toward 18:1n-9 (Fig. 5C) and not affect the Elovl5 activity toward 18:3n-6 (Fig. 5D). These results suggested that the GSK3-dependent phosphorylation of Elovl5 altered the substrate specificity of Elovl5 in C20 PUFA-treated cells.

### Phosphorylation of Elovl5 in the C-terminal region regulates the elongation activity toward 18:1n-9

To further support the involvement of the Elovl5 phosphorylation in the loss of elongation activity toward 18:1n-9, we mutated some of the predicted phosphorylation sites of Elovl5. N-terminal 3xFLAG-tagged wild-type Elovl5 (WT-Elovl5) and T281A/S283A/S285A triple mutant Elovl5 (3A-Elovl5) were separately transfected into HEK293T cells (Fig. 5E). The *in vitro* Elovl5 activity toward 18:1n-9 was increased in the cells transfected with WT-Elovl5 and 3A-Elovl5 (Fig. 5F). The increases in activities were correlated with the increases in protein expression levels. As expected, the Elovl5 activity in WT-Elovl5-transfected cells was almost completely suppressed upon addition of C20 PUFA, whereas the Elovl5 activity in 3A-Elovl5-transfected cells remained unaffected by C20 PUFA (Fig. 5F). The cellular level of 20:1n-9, a product of Elovl5, was also increased in the WT-Elovl5- and 3A-Elovl5-transfected cells, while in the presence of C20 PUFAs, it was greatly decreased in the WT-Elovl5-transfected cells but not in the 3A-Elovl5-transfected cells (Fig. 5G). Thus, phosphorylation of T281/S283/S285 appears to be necessary for the complete loss of Elovl5 activity toward 18:1n-9 in C20 PUFA-treated cells. The lower expression levels of 3A-Elovl5 protein than WT-Elovl5 (Fig. 5E) may support the idea that phosphorylation of Elovl5 protein confers its stability.

## Discussion

Concerning the synthesis of Mead acid in an EFA-deficient state, it has generally been assumed that Elovl5 has much higher activity towards 18:2 and 18:3 than 18:1n-9 (23), which results in efficient synthesis of arachidonic acid and EPA and that Elovl5 uses 18:1n-9 in the EFA-deficiency state simply because the preferred substrates (18:2 and 18:3) were absent. According to our study, this idea is partially true but insufficient. We showed that Elovl5 is constitutively phosphorylated and prefers 18:2 and 18:3 under normal nutritional conditions, but acquires an activity toward 18:1n-9 comparable to its activity toward 18:3n-6 by dephosphorylation of Elovl5 under the EFA-deficient conditions (Fig. 5H). This molecular mechanism should result in more precise regulation to Elovl5 for the synthesis of Mead acid as a substitute for arachidonic acid or EPA in an EFA-deficient state.

Mammalian elongases catalyze four separate enzymatic reactions (30), and it is generally believed that Elovls that catalyze the initial and rate-controlling condensation reaction between the fatty acyl-CoA and malonyl-CoA have different specificities for fatty acid substrates. Although seven Elovl isoforms with different substrate specificities have been identified to date, the molecular basis underlying the substrate specificity of each Elovl has not been fully elucidated. It has been elegantly demonstrated that Elo2, a yeast homologue of mammalian Elovls, determines the chain length of the end-product, but not one particular substrate, by utilizing the distance between the catalytic site (the HxxHH motif) and a lysine residue at the luminal surface of one transmembrane helix of Elo2 (31). In a study concerning the molecular basis for the differences between Elovl5 that can elongate C20 PUFA to C22 PUFA and Elovl2 that can elongate C20 PUFA to C22 PUFA and then to C24 PUFA (32), replacement of some amino acids in the 6th and 7th transmembrane domains of Elovl2 with the equivalent residues from Elvol5 resulted in a loss of the unique Elovl2 conversion of C22 PUFA to C24 PUFA, while retaining the ability to convert C20 PUFA to C22 PUFA. All these studies pointed out that the intrinsic structure of Elovls, especially that of the transmembrane domains, are critical for determining the substrate or the product specificity. The present study postulates a unique mechanism underlying the determination of the substrate specificity of mammalian Elovls: phosphorylation at the cytosolic C-terminus (not in the transmembrane helices) of Elovl5 abolishes the activity toward 18:1n-9 without affecting the activity toward 18:3n-6. Among fatty acid elongases, posttranslational modification has been reported only in yeast Elo2. Phosphorylation of Elo2 is increased in nutrient-depleted cells (26) or by inhibiting sphingolipid synthesis with myriocin, an inhibitor of serine palmitoyltransferase (27). Although it is controversial whether the phosphorylation of Elo2 enhances or suppresses its activity, phosphorylation of Elo2 appears to control the rate of VLCFA synthesis.

It is likely that phosphorylation/dephosphorylation of Elovl5 changes the substrate preference for 18:1n-9 based on the experiments using a GSK3 inhibitor and using the Elovl5 mutant substituting putative phosphorylation sites that are conserved in the yeast homolog Elo2. According to the Phos-tag experiments, Elovl5 appeared to be phosphorylated only partially under the C20 PUFA-supplemented conditions, under which it lost the activity towards 18:1n-9 almost completely. Recently, Elovl4, another fatty acid elongase with a sequence similarity to Elovl5, was shown to exist as a multisubunit complex, and that the complex is necessary for the functioning of Elovl4 as an elongase enzyme in the biosynthesis of VLCFAs (33). Assuming that Elovl5 forms a similar complex based on the sequence similarity and the common structural features of Elovls, phosphorylation of one or a few of Elovl5 in the complex might be sufficient for the shift of the substrate preference to 18:1n-9. Membrane topology studies have provided a model in which fatty acid elongases consist of a 7-transmembrane region in which the C-terminus faces the cytosol (31) and contains an intramembrane substrate-binding pocket (32, 34). Interestingly, amino acid residues that resemble the GSK3 consensus sites are critical for the alteration in substrate specificity and are located at the C-terminus. Thus, phosphorylation of the C-terminal amino acids may be involved in the change of the substrate preference to 18:1n-9. It remains unclear whether phosphorylation induces the conformational change near the substrate-binding pocket of Elovl5 or affects the interaction with an accessory protein(s) to the Elovl5 multisubunit complex in order to accomplish the change in substrate preference.

It also remains unclear how phosphorylation of Elovl5 was regulated in the EFA-deficient state. Phosphorylation of Elovl5 was abolished largely by BIO, a GSK3 inhibitor, but not by BIM, a protein kinase C inhibitor. However, it is unlikely that Elovl5 is directly phosphorylated by GSK3. GSK3 is a serine/threonine kinase that is constitutively active in its phosphorylated form, is primarily regulated by inhibition, and lies downstream of multiple cellular signaling pathways (35). GSK3 usually phosphorylates substrates that are pre-phosphorylated, and not substrates lacking priming phosphorylation. The Phos-tag experiments did not reveal a pre-phosphorylated form of Elovl5 in the presence of the GSK3 inhibitor, which suggests that the pre-phosphorylation of Elovl5 may not occur under C20 PUFA-supplemented conditions. So far, little is known about the intracellular signaling evoked by EFA-deficiency in mammalian cells. In yeast, when the VLCFA levels fall, Elo2 is controlled by signaling of the guanine nucleotide exchange factor Rom2, which is initiated at the plasma membrane (27). Low levels of VLCFA may limit the synthesis of sphingolipids in the plasma membrane, which regulates Rom2 activity toward a certain small G protein and the downstream signaling cascade for the Elo2 activation. A similar pathway leading to Elovl5 dephosphorylation might operate in mammals during EFA-deficiency. The present results show that the Elovl5 activity toward 18:1n-9 was controlled by 20:4n-6 and 20:5n-3, which are desaturated and elongated products of 18:2n-6 and 18:3n-3, respectively, but not by the EFAs 18:2n-6 and 18:3n-3 themselves. This indicates that more complicated mechanisms for sensing the membrane integrity than we had expected may initiate the signaling cascade for the Elovl5 phosphorylation/dephosphorylation. Alternatively, signaling lipid metabolites produced from 20:4n-6 and 20:5n-3 may control the pathways. Further studies are needed to determine how the cell senses the EFA-deficient state and how intracellular signaling evoked by C20 PUFAs leads to Elovl5 phosphorylation.

As mentioned earlier, Mead acid, in addition to substituting for essential PUFAs in biological membranes, has various biological activities and is associated with some diseases including cancer (16–19). A number of GSK3 inhibitors have been developed for the treatment of human diseases including type-II diabetes (36). In the present study, treatment with a GSK3 inhibitor induced the production of Mead acid even under normal nutrient conditions. Thus, in studies using a GSK3 inhibitor, caution might be needed to control the level of Mead acid because of its unusual bioactivity.

## Experimental procedures

### Materials

18:1n-9-CoA and 18:3n-6-CoA were obtained from Avanti Polar Lipids. Malonyl-CoA was obtained from Sigma-Aldrich. GSK3 inhibitor, 6-Bromoindirubin-3’-oxime (BIO) and PKC inhibitor, Bisindolylmaleimide I (BIM) were purchased from Cayman Chemical Company. Acetyl-CoA carboxylase inhibitor, 5-(tetradecyl oxy)-2-furoic acid (TOFA) was purchased from Santa Cruz Biotechnology.

### Cell culture and transfection

NIH3T3 and HEK293T cells were obtained from the American Type Culture Collection (ATCC), maintained in Dulbecco’s modified Eagle’s medium (high glucose) supplemented with 10% fetal bovine serum (FBS) and 100 U/ml penicillin, 100 mg/ml streptomycin, and 2 mM L-glutamine at 37 °C in a humidified atmosphere of 5% CO_2_. 48 h after seeding, NIH3T3 cells were treated for 24 h with a PUFA mixture [25 μM arachidonic acid (20:4n-6) and 25 μM EPA (20:5n-3)]. Transfections were performed using lipofectamine^®^ 3000 reagent (Invitrogen) according to the manufacturer’s instructions. Human Elovl5 was amplified by means of PCR assay from human liver cDNA, digested with BamHI, and inserted into the pCE-puro 3×1FLAG-1 vector. This vector was designed to produce an N-terminal 3×FLAG-tagged protein.

### siRNA transfection

Cells were plated in 6-well culture dishes at a density of 1.0 × 10^5^ cells per well. The cells were transfected using Lipofectamine^®^RNAiMAX reagent (Invitrogen) according to the manufacturer’s recommendations. A control siRNA and siRNAs directed against Elovl5 were purchased from Thermo Fisher Scientific.

### Total RNA isolation and quantitative real-time PCR

Total RNA was extracted from cells by using TRIzol Reagent (Invitrogen) and was reverse transcribed using a High Capacity cDNA Reverse Transcription kit (Applied Biosystems). Quantitative real-time PCR was preformed using SYBR Green PCR Master Mix and 7300 Real Time PCR System (Applied Biosystems). Primers were as follows: Elovl5 (fw, 5’-GGTGTGTGGGAAGGCAAATAC-3’; rv, 5’-TGCGAAGGATGAAGAAAAAGG-3’), Elovl6 (fw, 5’-TCAGCAAAGCACCCGAACTAGGTGA-3’; rv, 5’-ATGAACCAACCACCCCCAGCGA-3’), Elovl3 (fw, 5’-AGTGGGCCTCAAGCAAACCGTG-3’; rv, 5’-TGAGTGGACGCTTACGCAGGATGA-3’), Elovl7 (fw, 5’-AGCTCATGGAGAACCGGAAG-3’; rv 5’-GTACCCCAGCCAGACATCAC-3’). The relative target gene expression was normalized on the basis of GAPDH content.

### Fatty acid analysis by GC-MS

Cells were seeded in 6-well culture dishes at a density of 1.0 × 10^5^ cells per well and cultured for 72 h. Total cellular lipids were extracted using the method of Bligh and Dyer (37). Isolated lipids were methylated with 2.5% H_2_SO_4_ in methanol. The resulting fatty acid methyl esters were then extracted with hexane and quantified by gas chromatography–mass spectrometry (GC-MS) using a GC-MS QP2010 (Shimadzu) as described previously (13).

### in vitro fatty acid elongation assay

Cultured cells were suspended in buffer A [50 mM Hepes-NaOH (pH 6.8), 10% glycerol], and lysed by sonication. After ultracentrifugation (100,000 ×g, 30 min, 4 °C), the pellet was suspended in buffer A and was used as the total membrane fraction. Typical reaction mixtures of fatty acid elongation assays contained total membrane fractions (100 μg protein), 50 μM acyl-CoA complexed with 0.2 mg/ml FA-free BSA (Sigma), and 125 μM malonyl-CoA in a 250 μl reaction mixture (buffer A containing 150 mM NaCl, 2 mM MgCl_2_, 1 mM CaCl_2_, and 1 mM NADPH). After 30 min incubation at 37 °C, the reactions were terminated by adding 125 μl 75% KOH (wt/vol) and 250 μl ethanol, then saponified at 70 °C for 1 h. The reactions were stopped under acidic conditions with HCl. Lipid extract in hexane was dried, methylated, and then fatty acid methyl esters were quantified by GC-MS. The Elovl5 activity toward each fatty acid was determined by the amount increase of the elongated products.

### Western blot analysis

Equal amounts of protein were separated by electrophoresis on 10% SDS-PAGE. Proteins were transferred to Immobilon PVDF membranes and probed with rabbit anti-Elovl5 antibody, rabbit anti-Na^+^/K^+^ ATPase antibody (Cell signaling), mouse anti-beta-actin antibody (Abcam) and mouse anti-FLAG antibody (Wako). Anti-Elovl5 rabbit polyclonal antibody was produced with an immunogen corresponding to amino acids 255-267 of mouse Elovl5 (KKGASRRKDHLK). The secondary antibody was mouse anti-rabbit and goat anti-mouse IgG conjugated to horseradish peroxidase. Immunoreactive bands were visualized by enhanced chemiluminescence (GE Healthcare). Elovl5 phosphorylation was assessed by Phos-tag SDS-PAGE (38) followed by immunoblotting with Elovl5 antibody.

### Statistical analysis

Values represent means ± SEM from three independent experiments. Statistical analysis was performed by unpaired Student’s t-test or one-factor ANOVA of variance with a Tukey– Kramer post hoc test to identify significant differences.

## Acknowledgements

This work was supported by JSPS KAKENHI Grant Numbers JP17K00848. We are grateful to Makoto Murakami (University of Tokyo, Japan) for valuable comments on the study.

## Author contributions

Y.H., H.A., Y.F. and I.I. designed the study; Y.H., M.Y. and I.I. conducted the experiments; Y.H., M.Y., and I.I. analyzed the data; Y.H., N.K., H.A. and I.I. wrote the manuscript.

## The abbreviations used are

BIO: 6-bromoindirubin-3-oxime
EFA: essential fatty acid
Elo2: elongation of fatty acid protein 2
EPA: eicosapentaenoic acid
Elovl: elongase of very long chain fatty acid
FA: fatty acid
Fads: fatty acid desaturase
FBS: fetal bovine serum
GSK3: glycogen synthase kinase 3
PUFA: polyunsaturated fatty acid
TOFA: 5-(tetradecyloxy)-2-furancarboxylic acid
VLCFA: very long chain fatty acid

